# Detecting predicted cancer-testis antigens in proteomics datasets of healthy and tumoral samples

**DOI:** 10.1101/2024.06.08.597624

**Authors:** Karla Cristina Tabosa Machado, Tayná Da Silva Fiúza, Sandro José De Souza, Gustavo Antônio De Souza

## Abstract

Biomarkers are molecular markers found in clinical samples which may aid disease diagnosis or prognosis. High-throughput techniques allow prospecting for such signature molecules by comparing gene expression between normal and sick cells. Cancer-testis antigens (CTAs) are promising candidates for cancer biomarkers due to their limited expression to the testis in normal conditions versus their aberrant expression in various tumors. CTAs are routinely identified by transcriptomics, but a comprehensive characterization of their protein levels in different tissues is still necessary. Mass spectrometry-based proteomics allows the characterization of many cellular types and the production of large amounts of data while computational tools allow the comparison of multiple datasets, and together those may corroborate insights obtained at the transcriptomic level. Here a computational meta-analysis explores the CTAs protein abundance in the proteomic layer of healthy and tumor tissues. The combined datasets present the expression patterns of 17,200 unique proteins, including 241 known CTAs previously described at the transcriptomic level. Those were further ranked as significantly enriched in tumor tissues (22 proteins), exclusive to tumor tissues (42 proteins) or abundant in healthy tissues (32 proteins). This analysis illustrates the possibilities for tumor proteome characterization and the consequent identification of biomarker candidates and/or therapeutic targets.

## I. INTRODUCTION

Biomarkers are molecular components whose levels are altered in a given condition and whose presence in a sample allows its classification or indicates a clinical prognosis [1]. In the last decade, there has been an increase in the identification and characterization of such molecules in specific diseases [2]. Cancer-testis antigens (CTAs) are unique marker candidates for cancer diagnosis and prognosis due to their restricted expression [3]. They are normally present in healthy testis but are also aberrantly expressed in various tumor types, making them promising immunotherapeutic targets. Several clinical studies have been carried out or are in progress to explore the potential of CTAs as treatment targets in cancer [4]. For example, Li et al. performed RNA-seq analysis on glioblastoma samples and identified eight highly-expressed CTAs with the potential to predict tumor prognosis [5].

Biomarker candidates are initially identified through high-throughput omics techniques, by detecting differentially expressed genes when comparing normal and disease cells [6], [7]. Although transcriptomics is the most routinely used omics method, mRNA levels in a cell are not always proportional to protein levels for a given gene [8], [9]. Therefore, it is desirable that transcriptomics data is further characterized at protein level in healthy and disease tissues, to avoid false-positive predictions based solely on a single omics strategy [9], [10]. Proteomics has been regarded as a promising complementary technology that can provide insight into the disease at the protein level. [11]. It can provide protein abundance profiles of a given biological sample through the analysis of peptide mixtures using high resolution liquid chromatography coupled to mass spectrometry (LC-MS), where >10,000 proteins can be routinely identified [12]–[14].

The availability of raw and processed MS data has become routine for proteomics studies in recent years, and the construction of a consensus for MS data sharing allowed for the evaluation, reuse and comparative analysis of such data [15] and the consequent integration of information through meta-analysis techniques [16]. Meta-analysis is an efficient strategy for the integration and analysis of datasets from multiple studies and may provide more significant and reliable results [17], [18]. For example, the comparison of independently collected MS datasets allowed the integration of pancreatic cancer proteomic data to uncover proteins relevant to diagnosis and prognosis of pancreatic ductal adenocarcinoma [19], where the authors identified 39 secreted proteins with potential for serving as biomarkers. Rosenberg et al. performed a multivariate meta-analysis of proteomic datasets from prostate and colon tumors and identified 14 proteins with similar expression trend between healthy and tumor samples for both types, which were not found in the individual studies [20]. Shafi et al. developed a multicohort, multi-omics meta-analysis structure able to integrate independent studies centered around mRNA, DNA methylation, and protein-protein interactions, to identify biomarkers through a network analysis of glioblastomas and lowgrade gliomas. These biomarkers could explain the biological mechanisms of the diseases and predict survival of the patients [2].

In this study, we ran a computational meta-analysis on several proteomic datasets from tumor and healthy human tissues to verify protein levels of CTAs predicted by transcriptomics, and evaluate their potential as biomarker candidates. The integration of these datasets resulted in the total identification of 17,200 unique proteins and exploratory data analysis revealed that 241 of 418 CTAs previously predicted at the transcriptomic level were identified. Among these, 76 were more highly abundant in tumor tissues when compared to healthy tissue, while 32 were flagged as non-appropriate biomarkers due to high levels in healthy tissue samples other than testis.

## II. MATERIALS AND METHODS

### A. PROTEOMIC DATASETS

Approximately 10 Tb of raw mass spectrometry data were obtained from previously published works [12], [13], [14], [21], [22], [23], [24], [25], [26], [27], [28], [29], comprising more than 500 samples of healthy tissues, tumors collected from patients and immortalized cell line models. The following MS data were downloaded from ProteomeXchange: 11 human cell lines (PXD002395), 16 adenoma patient samples, 8 colon cancer tissues and their patient matched normal-appearing mucosa cells (PXD002137), the NCI-60 human tumor cell line panel (PXD005940, PXD005942, PXD005946), the HeLa proteome (PXD004452), ovarian cancer cell lines (PXD003668), subtypes of diffuse large B-cell lymphoma (PXD002052), 88 breast cancer clinical samples (PXD000815), breast cell lines and four primary breast tumors (PXD008222), four metastatic melanoma tissues (PXD001724), clinical samples from advanced stage melanoma patients (PXD006003), prostate cancer samples (PXD003430, PXD003452, PXD003515, PXD004132, PXD003615, PXD003636, PXD004159), healthy human heart (PXD006675), 29 healthy human tissues (PXD010154), and human immune cells (PXD004352). The above raw data were submitted to protein identification using MaxQuant version 1.5.2.8 [30], [31] using a non-redundant Uniprot human protein sequence database from July 2019 [32]. Parameters of selection include: mass error for intact peptides before calibration ≤ 20 pmm; mass error for intact peptides after calibration ≤ 6 ppm, error on the fragmentation spectra ≤ 0.5 Da; allowed amino acid modifications: oxidation of methionine (variable), N-terminal peptide acetylation (variable) and Cys carbamidomethylation (fixed); false-positive rate 0.01 (1%) calculated by reversed sequence strategy; maximum missed cleavage sites allowed = 2.

### B. DATA PROCESSING

The Maxquant’ `proteingroup.txt` output files were used for further analysis. These files contain information on the proteins identified from the raw data, as each row describes group of proteins that were reconstructed with a set of MS-identified peptides and includes, but are not limited to: the name of the genes expressing said group of proteins, total number of unique peptides associated to the group, number of proteins in group, and sample intensities of each protein group (which is the sum of intensities of peptides belonging to a group of proteins in a given sample).The pre-processing was implemented in R (v 4.0.5) and included removal of false-positives and contaminant identifications from each individual output file, integration of protein groups identifications into a single output, and transformation steps for data normalization, as shown below.

#### 1) Data cleaning

The script `processing_protein_groups.r` removed contaminants and false-positives from individual output files, as well as proteins with missing gene names. In addition, for protein groups containing more than one protein and one or more genes, `process_protein_groups.R` selected the protein entry number and its coding gene from the group member with the most number of supporting unique peptides. In rare occasions where the group contained more than one protein or gene names with equal number of unique peptides, the script kept only the first entry name reported by MaxQuant for simplicity. The *proteingroups*.*txt* files contain hundreds of attributes which are relevant to describe the quality and accuracy of the MS identification but are not essential for our analysis. To streamline further steps in our workflow, individual protein group files had some of those attributes removed, and only the protein identifier, gene name, number of peptides associated and protein intensity in the samples were kept.

#### 2) Data Integration

The resulting *proteingroups*.*txt* file structure was then modified, and the data was converted from a wide format to a long format through pivoting, to avoid the generation of a data frame with many missing values since not all files of groups of proteins contained the same individual proteins. Individual samples in each dataset were then classified and grouped according to their tissue of origin. The testis samples from the dataset of 29 tissues were not included in the Healthy grouping. The number of samples for each tissue of origin can be seen in Supplementary Data 1. Only tissues with at least ≥ 10 samples were considered, resulting in eight tumor types and one healthy tissue group. Finally, descriptive attributes for each sample were integrated to produce a data set containing: the database identifier, database name, tissue, sample id, sample name, gene name and protein intensity. Those steps were performed by the `integrating_protein_groups.R` script.

#### 3) Data transformation

To re-scale and normalize the intensity values for all proteins from the many independent studies, the script `normalizing_integrated_data.R` first transforms the intensity by a *log*_2_, then performs a Z-score normalization 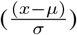. To replace missing values, we did a Random Tail Imputation (RTI) replacing missing points with random values from a normal distribution centered at the detection limit. This step was performed using the “rtruncnorm” function of the “truncnorm” R library. For proteins present in ≥ 5% of the samples, a specific data imputation approach was performed: random values drawn from a normal distribution where *µ* is the median of all intensity values for that protein and *σ* comes from the distribution itself. For proteins present in < 5% of the samples, imputation comprised random values drawn from a normal distribution where *µ* was given by the *µ* of the intensity values from all proteins minus 1.8 *\* σ*, and *σ* comes from the intensity values from all proteins *0.3 (*µ* = *µ*_*all*_ *− σ* * 1.8; *σ* = *σ*_*all*_ * 0.3) [33]. Imputation was carried out by the `imputing_integrated_data.R` script.

### C. PROTEIN SIGNATURE ANALYSIS

CTAs previously predicted at the transcriptomic level [34] were filtered from the group of identified proteins and their abundance in cancerous tissue and in healthy tissues were analyzed. To allow for a broad analysis of their patterns of occurrence, the values of median protein abundance among different samples of a given tissue type were categorized as: low (z-scores between -5.25 to-1.99); medium (-1.99 to 0.372); high (0.372 to 3.8); or absent, by the `converting_integrated_data.R` script. Such categorization was performed by an equal-frequency discretization method, which divided the dataset in ranges of values comprising the same number of data points. Each range was then labeled with one of the categories and the median abundance values were mapped accordingly [35]. Proteins ranked as medium or high in cancerous tissues, while low or absent in the healthy group, were selected. Statistical significance (p < 0.05) was calculated with a Student’s t-test using their z-score values for individual samples in each group. In parallel, proteins with medium or high median abundance in healthy tissue were also selected, regardless of their abundance in cancerous tissues. Their absolute abundance in each sample was then retrieved.

## III. RESULTS

### A. COMPUTATIONAL APPROACH AND DATA DESCRIPTION

A computational workflow was built to integrate and analyze independent proteomics datasets. The workflow starts with the analysis of raw mass spectrometry data using a non-redundant Uniprot protein sequence database, followed by the analysis of the protein identification files with R, covering data loading, cleaning, integration and transformation (Figure 1). Briefly, the public MS datasets of interest were compiled (Purple) and submitted to a peptide identification engine such as MaxQuant (Pink). The approach should be useful for outputs from any search engine, as long as the column header identifiers created by the engine are properly named in the individual post-search scripts. Search engine steps are not obligatory to be performed prior to our workflow, but it is important that the user guarantees that the independently obtained protein lists and the quantitative data from the independent datasets are comparable. For example, it is recommended that the protein lists use the same entry code format (same database source and version), to avoid misinterpretation during data integration steps. In this work, we had reanalyzed the datasets using MaxQuant, and all identified protein lists (proteingroups.txt) were loaded into the computation approach (Green). Shortly, the data is then processed to basically sort and filter relevant information in each dataset, compare identification lists while merging files, and normalize the protein quantitation across datasets (Blue). The final output is a single protein identification list containing all protein and gene identifiers found in all datasets, including the normalized quantitative information for each protein on each individual sample present in all datasets. This file can be then post-processed as required by the user.

**FIGURE 1.**
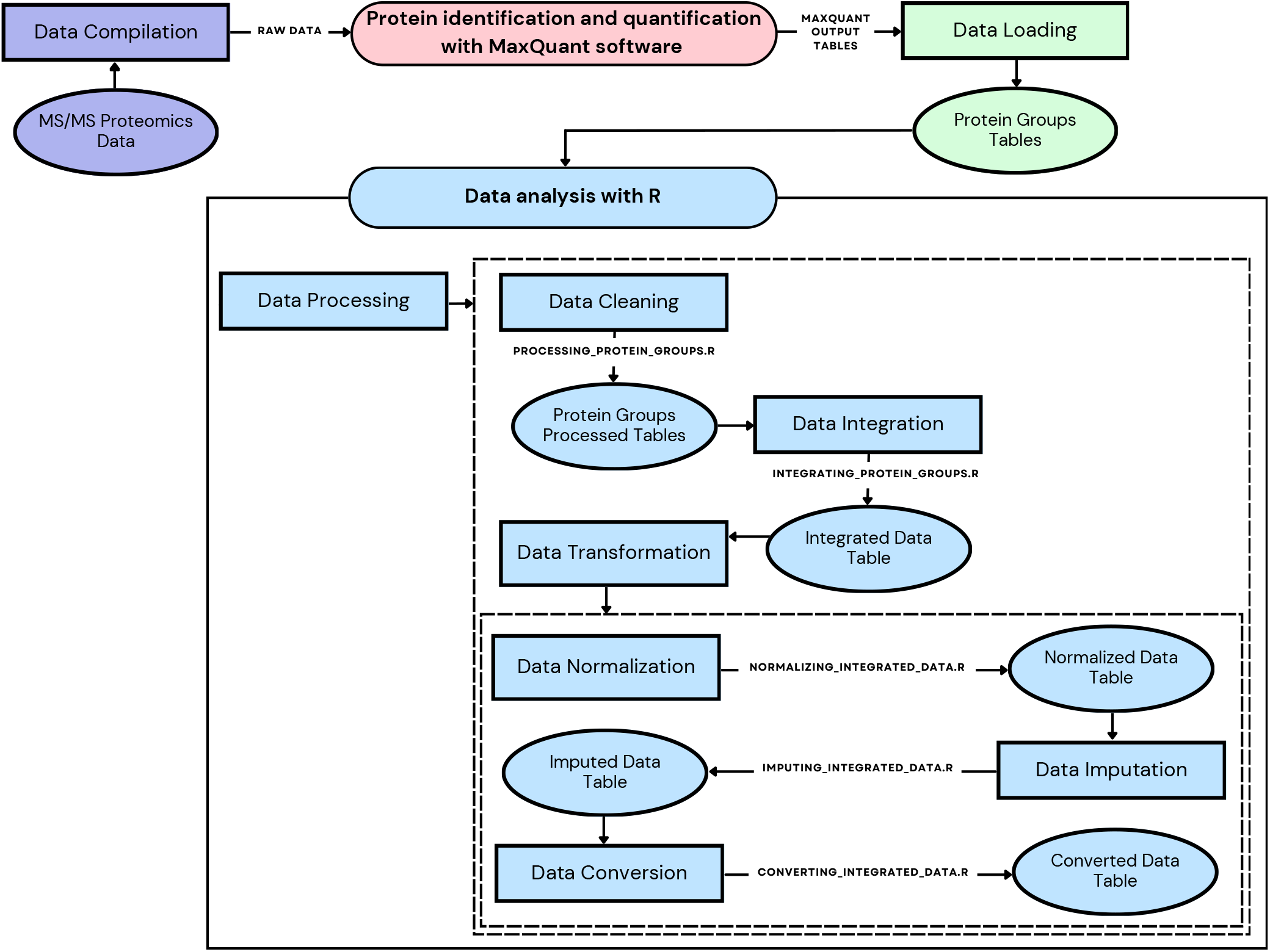
Overview of the meta-analysis of proteomic datasets. Publicly available raw mass spectrometry data were collected (Purple) and submitted to peptide identification using a MS search engine (Pink). For MaxQuant outputs, the resulting *proteingroups*.*txt* table files (Green) were then inputted into R data analysis. Files were processed as follow (Blue): data was cleaned (`processing_protein_groups.R`), removing contaminants and false-positive identifications, while also removing columns not relevant for post-processing; integrated (`integrating_protein_groups.R`) by finding shared and unique identifications across datasets; and transformed (`normalizing_integrated_data.R`, `imputing_integrated_data.R` and `converting_integrated_data.R`) to normalize quantitative information

### B. BIOMARKER CANDIDATES IN TUMORAL SAMPLES

Meta-analysis of the proteomic datasets identified 17,200 unique human proteins from 523 samples (138 from healthy tissues and 385 from various tumor tissues and immortalized cell lines). Individual samples per dataset were classified into categorical groups based on tissue origin, and proteins were then ranked as low/medium/high abundant based on their median distribution in each categorical group. As our research group had previously performed a large scale prediction of CTAs using public NGS data, we used this quantitative information to expand those findings at the protein level. The identified protein list was compared to 418 CTAs previously predicted at the transcriptomic level, from which 241 CTAs were observed at the protein level. Those 241 proteins and their respective categories for each tissue of origin can be seen in Supplementary Data 2. Their abundance profiles in all normal, healthy tissues plus testis are given in Supplementary Figure 1. Interestingly, 134 of those proteins were not identified at the protein level in testis. The remaining 107 proteins, 64 are either testis specific or dominantly abundant in testis, as expected from the previous mRNA prediction, while 43 proteins were observed with higher normalized quantitation in any other tissue than testis.

When also considering the abundance levels of those 241 CTAs in the cancer types analyzed, 76 had either medium or high abundance in at least one tumor type sample, while having low abundance (34 proteins, classified as “tumor enriched” from now on) or were not detected (42 proteins, classified as “tumor specific” from now on) in healthy tissues. Figure 2 shows a heatmap of those 76 CTAs allowing the visualization of the abundance patterns of those CTAs across cancer types that have greater expression in tumor tissues in comparison with healthy samples. From those, fifteen were classified as highly expressed in at least one cancer group, such as POTEF protein, highly present in B-cell lymphoma, lung, kidney and colon cancer samples, in addition of its medium abundance in breast cancer samples. This is an example of a tumor enriched identification, because some abundance was detected in tissues from the healthy group. BARHL2, which was only observed at higher levels in Prostate cancer samples, is an example of a tumor specific identification. Interestingly, just like BARHL2, half of the tumor specific CTAs (21 proteins) had a very tissue-specific molecular signature, being only observed in the prostate cancer samples.

**FIGURE 2.**
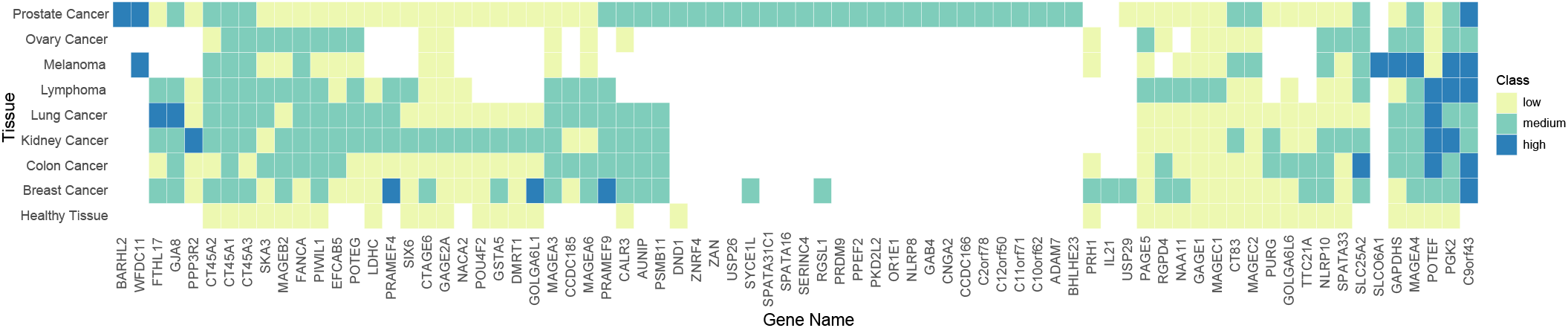
Heatmap of CTAs which are abundant in cancer samples when compared to healthy tissues. Pixel colors indicate low (light green •), medium (green •) or high abundance (blue •). White pixels indicate that the protein was not identified in any of the samples from the group. The abundance ranking was calculated from median z-score distribution for each individual protein per sample group

While class ranking is a good indication of possible marker signatures for each tumor tissue, a statistical analysis was performed in the tumor enriched group to verify if differences observed were significant while also considering intrasample variation. Z-score values of individual samples within each group were used, after data imputation for missing values. The statistical test showed that 22 of those 34 tumor enriched proteins were significant (p-values are given in Supplementary Data 3). Seven proteins were not significant (p-value *<* 0.05) and 5 proteins could not be subject to the t-test due to the low number of samples in which they were found. Such high percentage of significant proteins within those 34 CTAs indicates that the high/medium/low ranking performed is trustworthy. Figure 3 show examples of the sample quantitation distribution for the significant proteins GAPDHS and PGK2. GAPDHS was classified as highly present on melanoma, lowly on breast and prostate cancer and in B-cell Lymphoma, and with medium abundance in the remaining four cancer types (Figure 2). It was shown to be statistically significant in all of the investigated tumor samples with medium and high abundance. PGK2 was classified as medium presence in ovary and breast cancer samples, and highly present in the 6 other types of cancer, and was shown to be statistically significant in all cancer samples which it was detected.

**FIGURE 3.**
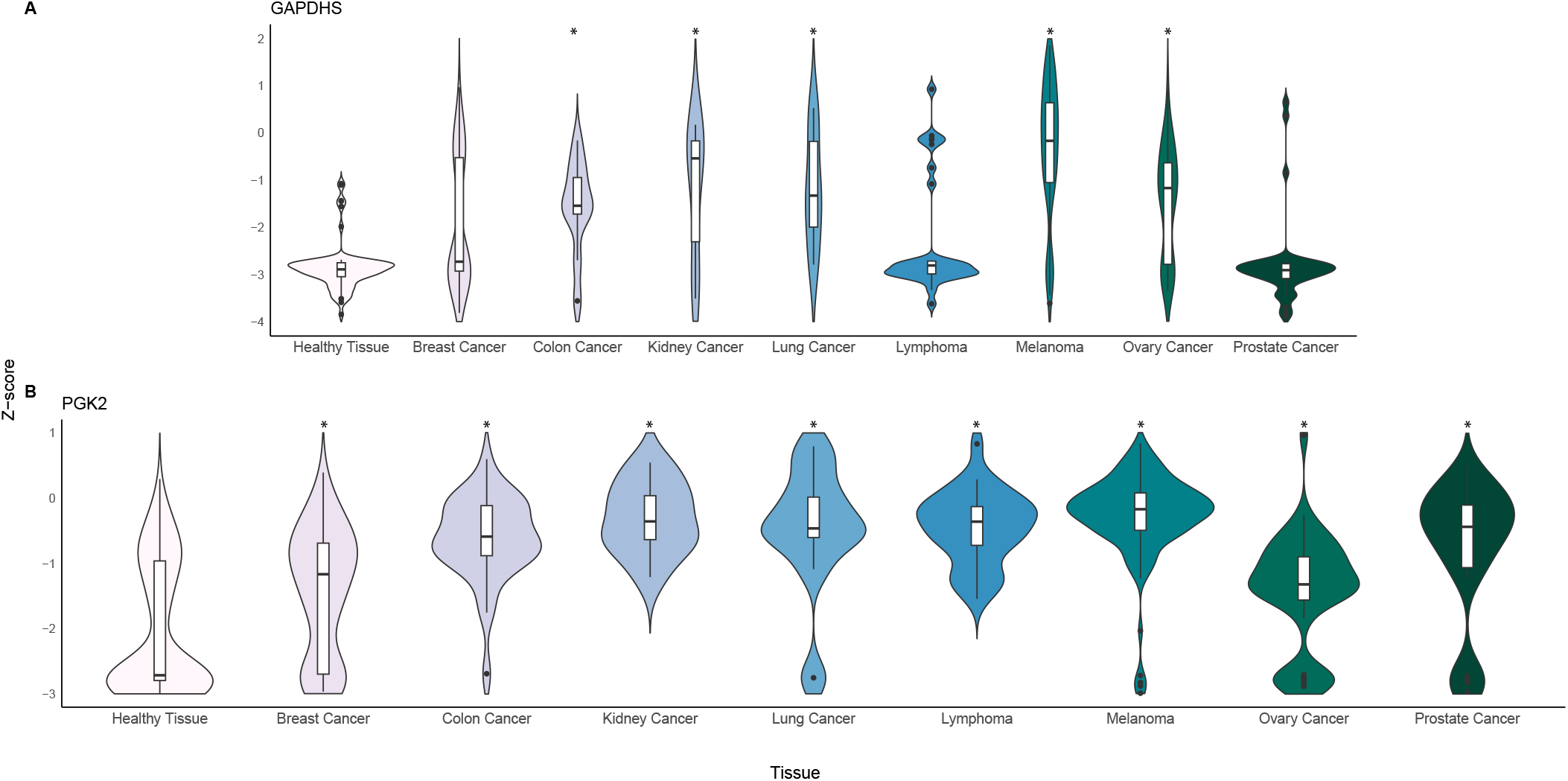
Quantitation distribution for proteins GAPDHS and PGK2 in all samples. The distributions are separated in the X-axis by the different cancerous tissues as well as the healthy group, while Y-axis shows individual Z-score values distribution for these proteins in each of the sample types. An asterisk indicates cancer groups for which there was a significant difference to the healthy group.

### C. DETECTION OF PREDICTED CTAS IN HEALTHY SAMPLES

Equally relevant are proteins that were predicted as CTAs at the transcriptomic level, but were detected with high or medium abundance in normal tissues at the protein level. Those CTAs were also selected regardless of their presence in tumor tissues (Figure 4). In total, 32 proteins predicted as abundant markers in cancer were found to have medium or higher abundance in healthy tissues. These identifications might also indicate current issues on CTA prediction based solely in transcriptomics. From those proteins, eight (KIF4B, SPATA31E1, SSX5, PCP2, OR2AK2, MAGEB4, GIP and LYZL2) had low or absent presence in any of the cancer groups under analysis. TUBA3C had the highest abundance measured in all groups, being highly present in 6 cancer types and in the healthy group. However, from the healthy group, TUBA3C was exclusively observed in samples from the normal heart dataset (Figure 5) [26]. Similarly, SPAM1, with medium abundance in prostate cancer, were only detected and quantified in the dataset with cells from the immune system.

**FIGURE 4.**
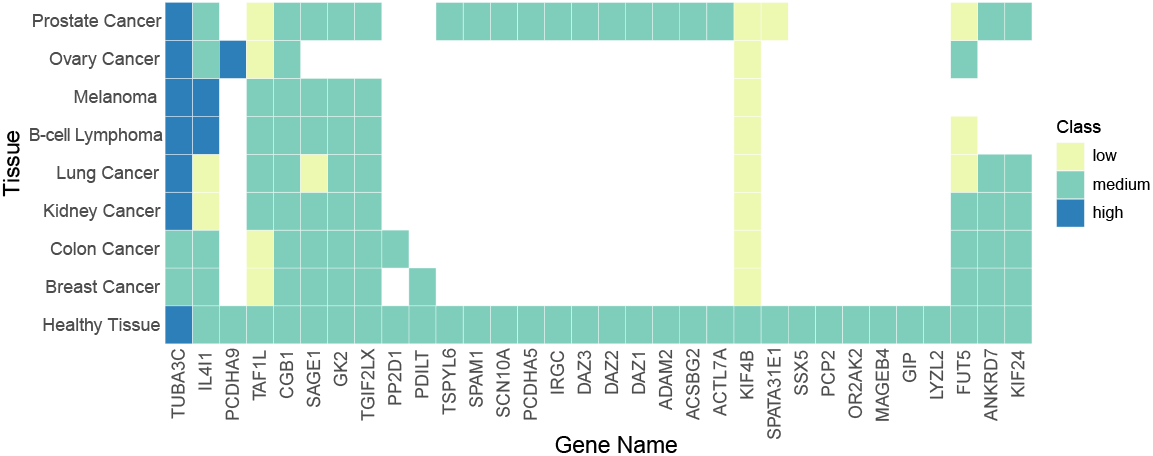
Predicted CTAs with high abundance in the healthy group. Proteins with high or medium abundance in the healthy group, regardless of their abundance on cancerous tissues, are shown. Colors indicate their category (low, medium or high) of abundance in each tissue. White squares indicate proteins absent in all samples of that group.

**FIGURE 5.**
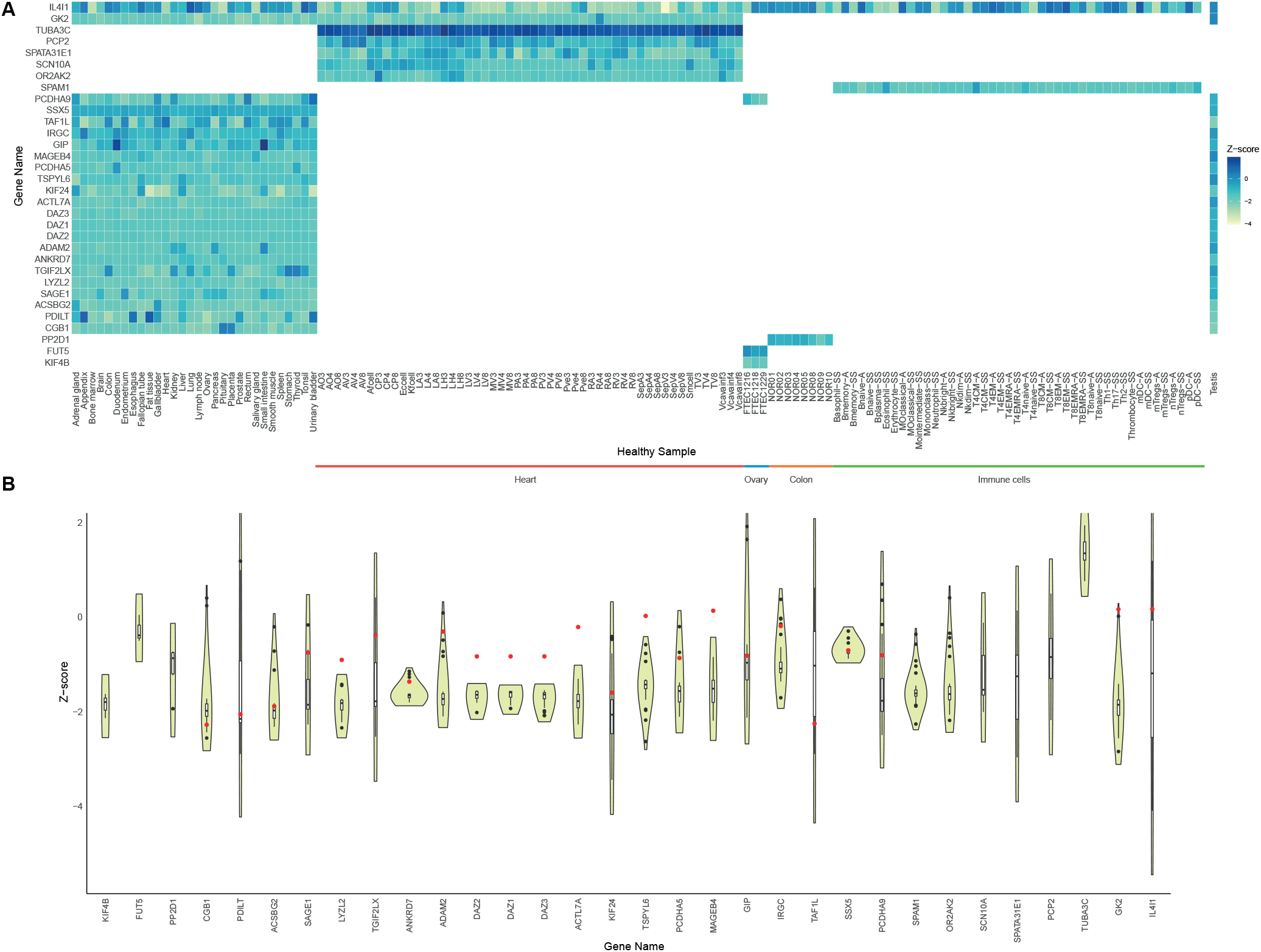
Individual quantitation of predicted CTAs which are abundant in the healthy samples. A: Heatmap of abundant CTAs in healthy tissues according to sample. Colors indicate the Z-score range for protein abundances in each sample. White represents that protein is absent in that sample. B: Z-score distribution of abundant CTAs in healthy samples, highlighting the testis measurement as a red dot compared to the distribution. The distributions are separated in the X-axis by gene name, while Y-axis shows Z-score values for these proteins in each sample healthy.

We also demonstrate the individual sample z-score contribution within the healthy group for those 32 proteins (Figure 5). Figure 5A represents the z-score values in every individual sample from the healthy group and Figure 5B shown a visualization of the distribution of the z-score value in healthy samples, contrasting with the z-score value of the testis (dot in red). Protein IL4I1 is the only in this set that was identified in all of the healthy samples tested, and its abundance is not dominant in the testis compared to the remaining healthy samples. TUBA3C, PCP2, SPATA31E1, SCN10A and OR2AK2 are observed only in cardiac tissue samples, while SPAM1 was only identified in cells from the immune system. Surprisingly, none of them were identified in testis (possibly absent or below MS detection capability). FUT5 and KIF4B are present in the fallopian tube (FTEC) samples while PP2D1 is present only in the colon tissue, and all of them are also absent in testis. The remaining identified CTAs are present in at least 29 different types of healthy tissues. Interestingly, only seven proteins (LYZL2, DAZ2, DAZ1, DAZ3, ACTL7A, TSPYL6, and MAGEB4) are dominantly present in testis, which was the criteria used in the transcriptomics approach to flag them as possible CTAs. But since their presence is also observed at significant levels in healthy tissues, these proteins should not be considered as specific antigen targets for cancer therapy.

## IV. DISCUSSION

The discovery of biomarkers for cancer often involves the selection of a group of candidate molecules based on their differential mRNA expression. Although it is a relevant approach, there is the risk that many predictions performed at the transcriptomic level do not correlate with protein abundance [8], [36]. The advance of proteomic technologies with the support of bioinformatic analysis has great potential in cancer research, allowing the identification and quantitation of proteins [37], [38] and, consequently, enabling the detection of biomarker candidates that may contribute to the diagnosis and/or personalized therapy [39]. Transcriptomic studies are still able to achieve a higher throughput result than that of proteomic experiments, but the integration of proteomic datasets to transcriptomic data enriches the body of information for the samples under study [40]. But proteomic studies demand a high expertise, specific instrumentation and experimental design and, most importantly, they are more time costly than NGS. So proteomic analysis is often aimed at identifying proteins in a specific experimental context, routinely comparing a limited number of samples. This demonstrates the relevance of developing broader, comparative, and quantitative proteomic meta-strategies, that can allow the further validation of transcriptomic datasets in areas such as in biomarker research [41].

Several studies have used integrative analyzes to investigate large volumes of proteomic and transcriptomic data, aiming to discover potential biomarkers for different types of cancers such as pancreatic cancer, breast cancer, and lung adenocarcinoma [19], [42], [43]. Such studies employ meta-analysis approaches and have identified targets that are potentially relevant for diagnosis, prognosis and even as therapeutic targets. But most importantly, meta-analysis in proteomics offers an opportunity to integrate multiple projects to interrogate predictions performed elsewhere. We selected 15 publications containing high protein identification coverage from public proteomic datasets of healthy and cancer samples. The data was integrated to further characterize the expression patterns of predicted CTAs previously identified by meta-analysis of transcriptomic data present in information derived from both the Human Body Map (HBM)/Genotype-Tissue Expression (GTEx)/Human Protein Atlas (HPA) and The Cancer Genome Atlas (TCGA). The complete proteomic meta-analysis of the selected proteomic studies resulted in the identification of 17,200 unique proteins. When we compared those with 418 previously identified CTAs in transcriptomic studies, 241 CTA candidates were also detected at the protein level.

Those identified CTAs were categorized into three groups where they seem to be: i) specific to one or more of the analyzed cancer samples, meaning high/medium abundance in that group and absent in healthy tissues or; ii) enriched in the cancer samples, meaning with high/medium abundance in cancer while low in healthy tissues or; iii) medium/high in healthy tissue regardless of their abundance in cancer samples. The majority of the enriched CTAs in tumor categories had individual sample abundance differences to be statistically significant, which strengthens the low/medium/high classification performed. The specific and enriched CTAs in cancer can be considered promising cancer markers for future studies. While CTAs that are specific in cancer may serve as therapeutic targets, the enriched CTAs likely do not have the same potential. This occurs because even with low abundance in healthy tissues, using these molecules as drug targets could also affect normal cells, resulting in undesired responses. But a more throughout investigation of healthy tissue protein abundance should be performed to confirm these findings.

A striking number of the predicted CTAs were invalidated when measuring its proteins levels. Only 64 were dominantly present in testis as expected, while 134 were not detected in in testis, 113 of those not detected in any healthy samples, including testis, and also had low detection in the analyzed cancer groups. This observation might be a result from methodological factors when comparing transcriptomic and proteomic datasets, but might also be due to the low exhautiveness of proteomic approaches. The 418 CTAs previously predicted at the transcriptomic level were identified using a proportional score, defined as the transcript level in a tissue divided by the sum of levels in all tissues, applying threshold of stringent 0.99, meaning that at least 99% of all expression of these genes in all analyzed tissues was derived from testis. Moreover, these genes had a level of expression (cutoff threshold of RSEM >1) in at least 15% of all informative samples for a given tumor [34]. This highlights the significance of proteomic analysis over solely relying on transcriptomic analysis, thereby strengthening the validation of the identified CTAs. For example, PCDHA9 had been previously identified as a CTA at the transcriptomic level because 99% of its total expression in the database was observed in testis, as described above. But in the proteomic datasets investigated here, only 2.59% of its total abundance was observed in testis. Those proteins should be then rejected as CTAs, since proteomic data indicates that their abundance distribution is more heterogeneous in healthy tissue group and not limited to testis at the protein level. However, from the 134 proteins not detected in either healthy samples or testis, they might still be valid antigen targets for other types of cancer that were not analyzed in the datasets selected in this work.

Some of the CTAs which protein detection were in agreement to RNA prediction had good correlation with known markers. For example, MAGEA4 is a highly studied CTA in melanoma cells [44], [45] and in triple-negative breast cancer (TNBC) [46]. CT45A1 shows significant over representation in breast cancer, and it has been demonstrated as a target for the development of new therapeutics [47], [48]. CT45A3 have a role in a specific ovarian cancer phenotype and is considered a prognostic factor [49]. CT45A2 is highly expressed in lung cancer and plays a protective role against this disease [50]. Our data confirmed all these observations, and also showed that MAGEA4 and CT45A1 are also enriched in all the remaining cancer groups investigated here.

Other tumor specific/enriched CTAs from our data are also potential biomarkers detected in other cancer types than the ones used in our study. AUNIP has been identified as a candidate biomarker in Oral Squamous Cell Carcinoma [51], pancreatic cancer [52], hepatocellular carcinoma and lung adenocarcinoma [53] and our data also suggests its potential relevance in breast, colon, kidney, lung, and prostate cancer. GJA8 was found to be overexpressed in Wilms tumor and might have a role an oncogene or tumor suppressor role [54]. Here, it exhibited significant levels in lung, breast, kidney, prostate and colon cancer, and in B-cell Lymphoma.

Finally, there are CTAs shown to be relevant in our dataset which to our knowledge are not reported in the literature. We observed high levels of SLC25A2 in colon cancer, and medium abundance in breast, kidney, ovary, prostate cancers, in B-cell lymphoma and melanoma. CDC185 was absent in healthy tissue and had medium abundance in colon and lung cancer, and B-cell lymphoma. WFDC11 also was absent in healthy tissue and high abundance in prostate cancer and melanoma. PSMB11 also was absent in healthy tissue and medium abundance in breast, colon, kidney, lung and prostate. Additionally, the roles of PRAME family members, such as PRAMEF4 and PRAMEF9, in cancer development are not well defined. This study suggests that PRAMEF4 might be relevant in breast, kidney, lung cancers and in B-cell lymphomas. PRAMEF9 might also be relevant in colon and prostate tissues.

In summary, our work demonstrates the importance of proteomic analysis as a complement to transcriptomic analysis to strengthen the validation of CTAs. This is because there is a risk that many predictions performed at the transcriptomic level do not correlate with protein abundance, which can lead to false-positive predictions when relying solely on a single omics strategy. Our data filtered a set of tumor specific/enriched CTAs from which roles should be further investigated for the tumor biology or its potential use for clinical applications. It also shows CTAs which are highly present in healthy samples and should therefore be classified as non-CTAs.

## V. CONCLUSION

This work presents a computational meta-analysis on several proteomic datasets from human tissues, both cancerous and healthy, aiming to identify at protein level the validity of CTAs predicted in transcriptomic repositories. This analysis highlights the potential of these proteins as candidates for cancer biomarkers or if their higher abundance in healthy tissues invalidate them for such. Identifying CTAs at the protein level is still evolving and there are still many challenges to be overcome, such as identifying low-abundance proteins in tumor cells. Our study has shown potential to use public proteomic datasets for the identification of candidate biomarkers, which may also serve as therapeutic targets.

## VI. DATA AVAILABILITY

All proteomic datasets collected and analysed in this study as well as code employed are available publically in: https://github.com/karlactm/metaanalysis and https://codeocean.com/capsule/1916652/tree.

## VII. AUTHOR CONTRIBUTIONS

KCTM: conceptualization, code implementations, methodology, composition and draft preparation; KCTM, TSF, SJS and GAS: composition, reviewing, and editing. GAS: conceptualization, supervision, funding, and infrastructure. All authors contributed to the article and approved the submitted version.

