## Supplementary figures and images for "Detecting predicted cancer-testis antigens in proteomics datasets of healthy and tumoral samples"

### Supplementary Figure 1

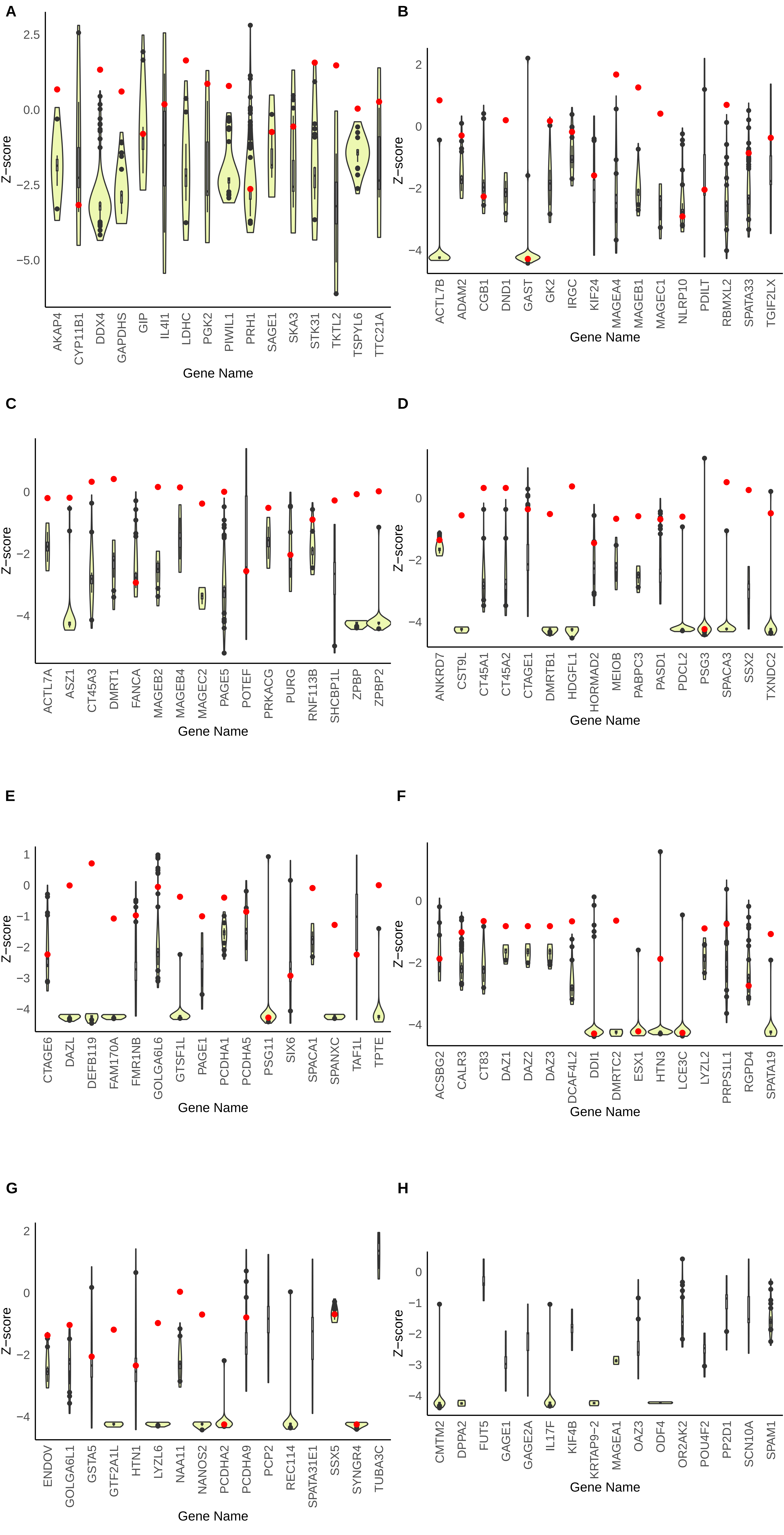
